# Hemispheric difference of adaptation lifetime in human auditory cortex measured with MEG

**DOI:** 10.1101/2024.04.16.588236

**Authors:** Asim H. Dar, Nina Härtwich, Aida Hajizadeh, Michael Brosch, Reinhard König, Patrick J. C. May

**Affiliations:** Leibniz Institute for Neurobiology, Research Group Comparative Neuroscience, Magdeburg, Germany; Fraunhofer Institute for Factory Operation and Automation IFF, Magdeburg, Germany; Department of Psychology, Lancaster University, Lancaster, UK

## Abstract

Adaptation is the attenuation of a neuronal response when a stimulus is repeatedly presented. The phenomenon has been linked to sensory memory, but its exact neuronal mechanisms are under debate. One defining feature of adaptation is its lifetime, that is, the timespan over which the attenuating effect of previous stimulation persists. This can be revealed by varying the stimulus-onset interval (SOI) of the repeated stimulus. As SOI is increased, the peak amplitude of the response grows before saturating at large SOIs. The rate of this growth can be quantified and used as an estimate of adaptation lifetime. Here, we studied whether adaptation lifetime varies across the left and the right auditory cortex of the human brain. Event-related fields of whole-head magnetoencephalograms (MEG) were measured in 14 subjects during binaural presentation of pure tone stimuli. To make statistical inferences on the single-subject level, additional event-related fields were generated by resampling the original single-trial data via bootstrapping. For each hemisphere and SOI, the peak amplitude of the N1m response was then derived from both original and bootstrap-based data sets. Finally, the N1m peak amplitudes we used for deriving subject-and hemisphere-specific estimates of adaptation lifetime. Comparing subject-specific adaptation lifetime across hemispheres, we found a significant difference, with longer adaptation lifetimes in the left than in the right auditory cortex (*p* = 0.004). This difference might have a functional relevance in the context of temporal binding of auditory stimuli, leading to larger integration time windows in the left than in the right hemisphere.

## 1 Introduction

The auditory system exhibits a plethora of hemispheric lateralisations in both structure and function (for a review, see Ruthig & Schönwiesner, 2022; Ocklenburg & Güntürkün, 2024). In humans, language processing serves as a prime example, as it was originally thought to reside predominantly in the left hemisphere (Broca, 1861; Wernicke, 1874). Evidence from brain-imaging, lesion, and intracortical studies point to the left hemisphere tracking temporal information needed for deciphering the rapidly changing nature of speech over time, whereas the right hemisphere shows specialisation in processing spectral information such as pitch (Cha et al., 2016; Schönwiesner et al., 2005; Zatorre et al., 2002). Further, there are asymmetries in brain activation associated with the processing of non-speech sounds: First, the temporal lobe of the left hemisphere appears to handle the categorisation of acoustically complex non-speech sounds, suggesting that this region supports new, task-required expertise in auditory processing (Leech et al., 2009). Second, categorisation of the pitch direction of frequency-modulated sounds preferentially activates the posterior auditory cortex (AC) of the left hemisphere, whereas categorisation of the duration of these sounds elicits stronger activity in the right posterior AC (Brechmann & Scheich, 2005). Thus, the hemispheric asymmetry in the processing of sounds is specific to both task and stimulus. Given the accumulated evidence on hemispheric differences, we investigate in this study whether neuronal processes related to auditory sensory memory also show lateralisation. To do this, we exploit a classic index of memory in sensory processing, that is, the phenomenon of adaptation: the diminishing of neuronal responses with stimulus repetition.

Adaptation, also called repetition suppression, is observed in all parts of the auditory system (Willmore & King, 2023). The effect can be characterised through its magnitude—by how much the neuronal response diminishes—and through its lifetime, the timespan over which the attenuating effect of previous stimulation persists. In general, adaptation lifetime is shorter in subcortical stations (tens of milliseconds) than in AC (seconds), and it is also longer in the non-lemniscal pathway than in the lemniscal one (Parras et al., 2017; Pérez-González & Malmierca, 2014). Adaptation indicates that previous stimuli leave an imprint in the brain that affects the processing of incoming stimuli. As these imprints encode the past stimulation, they can be regarded as sensory memory traces.

In humans, adaptation can be observed non-invasively in the event-related field (ERF) of the magnetoencephalogram (MEG) and in the event-related potential (ERP) of the electroencephalogram (EEG). In the auditory domain, the N1m response of the ERF and its electric counterpart, the N1, exhibit strong adaptation (for reviews, see May & Tiitinen, 2010; Näätänen & Picton, 1987). The N1m/N1 is a prominent wave of the auditory event-related response, peaking at approximately 100 ms after stimulus onset, and represents widespread activation of AC. Using MEG, Lu and colleagues (1992) found that the adaptation lifetime of the N1m of a subject correlates with the duration of their auditory sensory memory in a behavioural forced-choice comparison task for loudness. Further, the adaptation of the N1m/N1 response exhibits stimulus specificity, which is reflected in the oddball stimulation paradigm, when often-repeating sounds (standards) are occasionally replaced by a deviating sound. The difference in magnitude between these two responses is termed the mismatch negativity (MMN). The MMN shows that adaptation does not generalise to all stimuli and is instead specific to the standard stimulus, whereas the responses to the deviants tend to be of higher magnitude, as originally demonstrated by Butler (1968). The MMN is the traditional non-invasively measured index of sensory memory because a differential response to an incoming deviant indicates that the recent sensory past comprising the repetitive standard must somehow be represented in memory (Näätänen et al., 1978). Importantly, evidence shows that MMN is linked to stimulus detection and discrimination: First, the latency of the MMN predicts behavioural reaction time to the deviant (Tiitinen et al., 1994) and, second, the presence of the MMN predicts the ability of the subject to detect the deviant (Näätänen et al., 1993).

The above results suggest that the stimulus-specific adaptation taking place in AC, as observed in N1m/N1 and MMN responses, can be described as a behaviourally consequential memory effect. However, an alternative interpretation of these responses, not based on memory, is expressed in the predictive coding framework (Friston, 2005; Rao & Ballard, 1999; Wacongne et al., 2012). According to this view, the N1m/N1 adaptation is due to a suppressive top-down prediction signal and the MMN signifies a prediction error signal (for a critical review, see May, 2021).

Previous ERF studies have looked at the adaptation lifetime *τ*_SOI_ of the N1m response in the two hemispheres. Lu et al. (1992) and Rojas et al. (1999) presented tones to either the left or the right ear and found no hemispheric differences of the adaptation lifetime. McEvoy et al. (1997) presented tones and tone pairs to the left ear. Again, there were no statistically significant differences in *τ*_SOI_ between the left and right hemisphere. Zacharias et al. (2012) used monaural stimulation to the left ear and found that the mean *τ*_SOI_ was larger in the left hemisphere (2.8 s) than in the right (2.2 s); however, no statistical analyses were carried out. Cheng & Lin (2012) studied the effect of aging on adaptation lifetime using binaural stimulation with a train of five tones. Five different trains were presented, each characterised by a constant SOI between 0.5 s and 8 s. They report a gradual decrease in adaptation lifetime from younger to elderly subjects, but no hemispheric differences.

In summary, there is no conclusive picture on the lateralisation of adaptation lifetime. The majority of previously conducted studies utilised monaural or alternating binaural stimulation, and thus more strongly activated the contralateral cortex. This means it remains unclear whether there are inherent differences in adaptation lifetime across the hemispheres in the more natural situation of binaural stimulation. Moreover, previous studies did not report *subject-specific* statistics regarding the lifetime of adaptation in each of the hemispheres. In our MEG study, we developed a new analysis pipeline, utilising the bootstrap technique, to compute such subject-specific statistics and explored whether adaptation lifetime shows hemispheric differences. We presented binaural pure tones in separate blocks that differed in SOI. From the corresponding ERFs, the adaptation lifetime was then estimated individually for each subject from the peak value of the N1m response in the left and right hemisphere.

## 2 Methods

### 2.1 Subjects

Fourteen healthy participants (11 males, 3 females, mean age 33.8 years) with normal audiograms contributed to the MEG data of this study, which was approved by the Ethics Committee of the Otto-von-Guericke University Magdeburg. The subjects mainly belonged to the academic environment of the Leibniz Institute for Neurobiology, Magdeburg and the Otto von Guericke University, Magdeburg. All subjects agreed in writing to participate in the MEG measurements.

### 2.2 Stimuli and experimental design

A regular-SOI paradigm was used for the MEG measurements, which consisted of ten blocks, each comprising a sequence of 1.5-kHz tones of 100-ms duration including a linear rise and fall time of 5 ms, respectively. In each block, the tones were separated by a constant SOI, which was characteristic for this block. To improve the signal-to-noise ratio for short SOIs, tones presented in blocks with SOIs of 0.25 s, 0.5 s, and 0.75 s were repeated 120 times. In order to keep the total duration of the measurement at a comfortable level for the subjects, tones presented in the blocks with SOIs of 1 s, 1.5 s, 2 s, 3 s, 4 s, 5 s, and 7 s were repeated only 100 times. A full measurement session took approximately 45 minutes. The sequence of blocks was randomised across subjects.

The tones were generated with a conventional PC using the commercial *Presentation* software package (Neurobehavioral Systems Inc., Albany, CA). They were delivered to both ears of the subjects, simultaneously, via a plastic tube that terminated in an earmold. The tone presentation for each block was individually set off by the subject pressing a distinct button of a keypad. Between consecutive blocks, participants were allowed short breaks, which typically varied between about 10 s to 30 s. For each subject, the sound pressure level (SPL) was adjusted to 80 dB SPL for each ear before the measurement, with two exceptions where the SPL in the left ear was set to 75 dB at the request of the participants (subjects ‘i’ and ‘n’). All subjects reported the same subjective level of loudness in both ears. Further, subjects were instructed not to move and to focus on a fixation cross that was projected on a screen 1 m in front of them.

### 2.3 Data Acquisition and Analysis

#### 2.3.1 Acquisition and pre-processing

All MEG measurements were recorded at a sampling rate of 1000 Hz using a whole-head Elekta Neuromag TRIUX MEG system. The MEG device was located in a magnetically shielded and ventilated chamber containing a camera and loudspeakers so that the experimenter could visually observe and communicate with the subject. During data acquisition, an online filter was used with a low-frequency cut-off of either DC or 0.1 Hz, a high-frequency cut-off of 330 Hz, and the signal-space separation (SSS) method was applied to the data. To detect artefacts caused by eye movements and eye blinks, concurrent measurements of horizontal and vertical electrooculograms were taken.

Raw data from all 102 magnetometers was loaded into the Brainstorm software (Tadel et al., 2011) for pre-processing. Trials contaminated by eye blinks were rejected from all MEG channels. We used the signal space projection (SSP) module included in Brainstorm for the detection and correction of heartbeat artefacts in subjects whose MEG signal showed prominent regular heartbeat cycles. Further, data was visually inspected for any other artefacts such as muscle tension or technical incidents in the signal. After pre-processing, continuous data from all channels was segmented into epochs ranging from -500 ms to 1500 ms, with stimulus onset at time *t* = 0. All artefact-containing epochs were discarded. Furthermore, the non-adapted response to the first stimulus in each block was omitted (Rosburg et al., 2010). The remaining epochs (depending on SOI, these were typically 80 – 90 % of all epochs) were exported for further analysis within the Julia language (Bezanson et al., 2017). The exported data were band-pass filtered (high-pass 1 Hz, low-pass 30 Hz, both zero phase, Butterworth design of order 5).

#### 2.3.2 Event-related fields and their dependence on SOI

For each SOI block, ERFs were computed by arithmetically averaging all artefact-free epochs of single-trial activity. In our default analysis, all ERFs were baseline corrected based on the 200-ms interval prior to stimulus onset. However, to verify the baseline-corrected findings, we also ran the complete analysis procedure without baseline correction. Since ERFs with substantial amplitudes were obtained exclusively in MEG channels located above the AC of the left and the right hemisphere, we constrained the analysis to two respective subsets of channels, one for each hemisphere. The posterior MEG channel of each hemisphere that for most SOIs showed the largest absolute N1m-peak amplitude in the time window between 50 ms and 150 ms was chosen as the *principal channel*. All subsequent analysis steps were performed on data recorded by this principal channel. Figures 1a, b show, as an example, the SOI-dependence of the ERFs of a single subject recorded with the principal MEG channel above the left and the right hemisphere, respectively.

**Figure 1:**
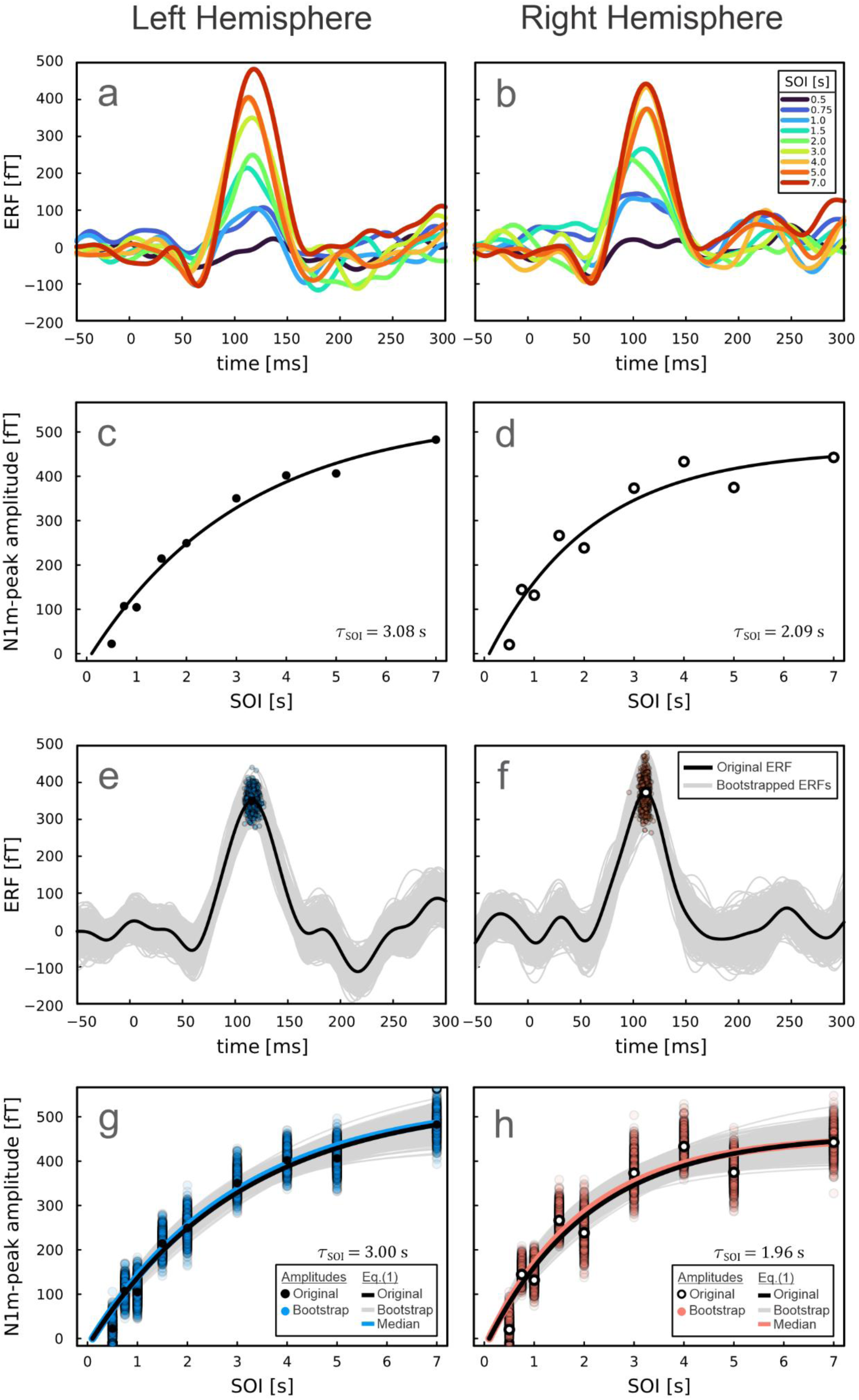
Procedure for determining the adaptation lifetime *τ*_SOI_ using the example of a single subject. Panels in the left column refer to the left hemisphere; those in the right to the right hemisphere. **a, b)** SOI-dependence of ERFs of this subject. The individual waveforms represent ERFs from SOIs between 0.5 s and 7 s, recorded by the subject-and hemisphere-specific principal MEG channel. With increasing SOI, the N1m-peak amplitude gradually increases and converges towards a saturation value at the largest SOIs. Since the magnetic field patterns of auditory ERFs have different polarities in the two hemispheres, the ERFs in b) were multiplied with -1 to enable an easy comparison between the two hemispheres. Stimulus onset is at time *t* = 0. **c, d)** Derivation of *τ*_SOI_ from the SOI-dependence of the N1m-peak amplitudes of the data shown in a) and b) for the left (closed symbols) and the right hemisphere (open symbols). The curves are fits of Eq. (1). **e, f)** Generation of surrogate ERFs using the bootstrap technique. ERFs for the 3-s SOI from each hemisphere are displayed as an example. The black curves represent the original ERF waveforms. The light grey waveforms represent the 999 bootstrap-based ERFs for each hemisphere, with the corresponding N1m-peak amplitudes displayed as blue (e) and pink (f) circles. **g, h)** Fits of Eq. (1) to the SOI-dependence of N1m-peak amplitudes from the original (closed and open black circles, see (c) and (d), respectively) and the resampled data (blue and pink circles, see (e) and (f), respectively). The fits to the resampled data are shown as light grey curves. The blue (left hemisphere) and pink (right hemisphere) curves depict Eq. (1) computed with the median values of τ_SOI_ and *A* across all fits.

For quality assurance of the peak amplitude of the N1m, we determined a signal-to-noise ratio (SNR) for each subject, hemisphere, and SOI. The SNR was defined as the ratio between the N1m-peak amplitude and the standard deviation of the MEG signal in the pre-stimulus interval of −200 ms to 0 ms. Based on a review of all ERFs and the associated SNR-values, we defined an SNR threshold of 1.5. Next, ERFs with an SNR < 1.5 that lacked a distinct N1m peak were excluded. This led to the exclusion of most of the 250-ms-SOI ERFs. In consequence, we decided to remove all 250-ms-SOI ERFs from further analysis to ensure fairer comparisons across the entire cohort. Furthermore, to keep the number of SOIs used for the estimation of adaptation lifetime the same for the two hemispheres of a given subject, data from *both* hemispheres was removed from further analysis for a given SOI, even if only one MEG channel, either in the left or in the right hemisphere, had an SNR value below this threshold.

#### 2.3.3 Deriving the adaptation time constant

The SOI-dependence of the N1m-peak amplitudes of ERFs, as shown in Figures 1a and b, enables the estimation of adaptation lifetime. Figures 1c and d illustrate, as an example, the N1m-peak amplitudes from Figures 1a and b as a function of SOI for the left hemisphere (closed symbols) and the right hemisphere (open symbols). In the standard regular-SOI paradigm, the peak amplitude of the N1m response grows as a function of SOI. This happens at first rapidly when SOI < 2 s, and then more slowly as peak amplitude levels off at longer SOIs, converging towards a maximum. This relationship can be well approximated by a single saturating exponential function (Hajizadeh et al., 2022; Lu et al., 1992):

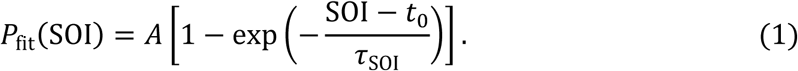

The time constant *τ*_SOI_ determines the rate of growth of the function towards the saturation amplitude *A* and is an estimate of adaptation lifetime. The term *t*_0_ determines the intercept of the function with the abscissa.

In our analysis, *t*_0_ was set to the tone duration, i.e., *t*_0_ = 100 ms. The reasoning behind this was that for *t*_0_-values shorter than tone duration, individual tones would no longer be distinguishable from one another. We applied the least squares method (LsqFit.jl, Julia) with *A* and *τ*_SOI_ as free parameters to fit Eq. (1) to our data. The results of the fits for the two hemispheres of one subject are indicated by the black curves in Figures 1c and d, respectively. To ensure robust fitting results for *τ*_SOI_, we applied a two-step approach: First, we determined a *τ*_SOI_-value for each subject and hemisphere by using an arbitrary starting value of *τ*_SOI_ = 0.1 s for the fitting procedure. Next, we computed the arithmetic mean of these *τ*_SOI_-values across all subjects and hemispheres and used this as the starting value (*τ*_SOI_ = 1.3 s) for the second iteration. In both steps, the starting value for *A* was the maximum peak-amplitude value across all SOIs, separately for each hemisphere and subject. For further verification, the entire analysis was also repeated with a three-parameter fit, where *t*_0_ was set as a free parameter with a lower bound of 100 ms and an initial value of *t*_0_ = 100 ms.

The small sample size (here: S = 9 SOIs) available for the fit of Eq. (1) to the N1m-peak amplitudes may raise concerns about the quality of the estimates of the fitting parameters. Hence, to gauge the robustness of our results, we generated surrogate waveforms at each SOI by applying the non-parametric bootstrap technique (Efron, 1979; Sielużycki et al., 2021). This approach yields information about statistical inferences such as median or mean and confidence intervals (CIs) without the restriction that the data are normally distributed or homoscedastic. We computed multiple realisations of the ERF waveforms for each subject and hemisphere by resampling trials according to the following procedure. For each SOI block, trials were randomly drawn with replacement from the pool of artefact-free trials to generate bootstrapped ERFs of the same dimensions (number of time samples and total number of trials) as the original trial-averaged ERF. This procedure was repeated to generate 999 bootstrapped ERFs, thus providing 1000 ERFs including the original, for all SOI blocks that met the criteria for acceptance (see Sect. 2.3.2).

Figures 1e, f show examples of this approach for the 3-s-SOI block. The black curves represent the original ERFs, as depicted in Figures 1a and b, and the closed black (Figure 1e) and open black (Figure 1f) symbols denote the N1m-peak amplitude from the respective original data for the left and right hemisphere, as depicted in Figures 1c and d. The light grey curves represent the bootstrap-generated ERFs, with their respective N1m peaks highlighted as blue (left hemisphere, Figure 1e) and pink (right hemisphere, Figure 1f) circles. The grey band mapped out by the bootstrapped ERFs illustrates the variability across the single-trial responses that form the basis of the original ERF. Similarly, the scatter of the associated N1m-peak amplitudes provides an estimate of the different CIs (for example the 95-% CI) associated with the original peak amplitude.

In the final step of the analysis, all N1m-peak amplitudes (i.e. 1000 for each SOI) were used to generate 1000 sets of data, each set containing an N1m-peak amplitude for each SOI. The sampling of the peak amplitudes from the bootstrapped pool to generate these data sets was done randomly and without replacement. Eq. (1) was fitted to each of the 1000 sets separately, thus obtaining 1000 individual exponential curves. Figures 1g, h show, for each hemisphere, the original data (closed and open black symbols) and the fit of Eq. (1) to these data (black curve). The vertical distribution of blue and pink symbols at each SOI are the N1m-peak amplitudes from the bootstrapped ERFs (see Figures 1e, f) and the grey curves represent the fits to the described sets of this data. The grey band mapped out by the bootstrap-based fits reflects the range of fitting parameter values resulting from the variability across the single-trial responses and thus provides CIs for the *τ*_SOI_-and *A*-values associated with the original ERFs. From the parameter values for all fits, we extracted the median values for *τ*_SOI_ and *A*. These two medians, together with the fixed *t*_0_ = 100 ms, were entered into Eq. (1) to compute the final response recovery function. For the illustrative data shown in Figures 1g and h, we obtained *τ*_SOI,LH_ = 3.00 s and *τ*_SOI,RH_ = 1.96 s for the adaptation lifetimes, and *A*_LH_ = 539 fT and *A*_RH_ = 456 fT for the saturation amplitudes, respectively (subscript LH corresponds to left hemisphere and RH to right hemisphere).

## 3 Results

### 3.1 Adaptation lifetime across subjects and hemispheres

Figure 2 shows the SOI-dependence of the peak amplitude of the N1m separately for each of the fourteen subjects. Black solid symbols refer to the original data from the left hemisphere; black open symbols refer to those from the right hemisphere. Blue and pink curves represent the exponential function of Eq. (1) computed with the respective bootstrap-derived median τ_SOI_-and *A*-values (see Methods). The corresponding median hemisphere-specific τ_SOI_-values are displayed in each panel. A comparison of the adaptation lifetime τ_SOI_ derived for the left and the right hemisphere of each subject is shown in Figure 3a, along with the corresponding 95-% CIs. Median τ_SOI_-values cover a range from 1.0 s to 3.0 s in the left and from 0.4 s to 2.0 s in the right hemisphere, and the corresponding CIs extend from 0.8 s to 4.9 s in the left and from 0.2 s to 2.9 s in the right hemisphere. We quantified the relationship between 95-%-CI ranges and respective medians via the ratio (*q*_97.5_(*x*) − *q*_2.5_(*x*))⁄*m*(*x*), where *x* denotes one subject-and hemisphere-specific distribution of *τ*_SOI_-values, *q*(*x*) the respective quantiles, and *m*(*x*) the median of the distribution. Across subjects and hemispheres, this ratio ranged from 0.34 to 1.21, with a grand median of 0.66. Thus, overall, 95-% CI ranges were large relative to the respective median *τ*_SOI_-values. We found that the median τ_SOI_-value for the left hemisphere was larger than that for the right hemisphere in the majority of subjects (12 out of 14). A non-parametric Wilcoxon signed rank test confirmed that this difference is statistically significant (estimated z = 2.73, two-sided *p*-value = 0.004) and that the effect size r = *z*⁄√*n* = 0.73 is large (see, for example, Fritz et al., 2012). Note that, since the τ_SOI_-values for the left and right hemisphere are subject-specific, comparing hemisphere-specific grand means of τ_SOI_ is not as meaningful and the hemisphere-specific values should be compared for each subject individually.

**Figure 2:**
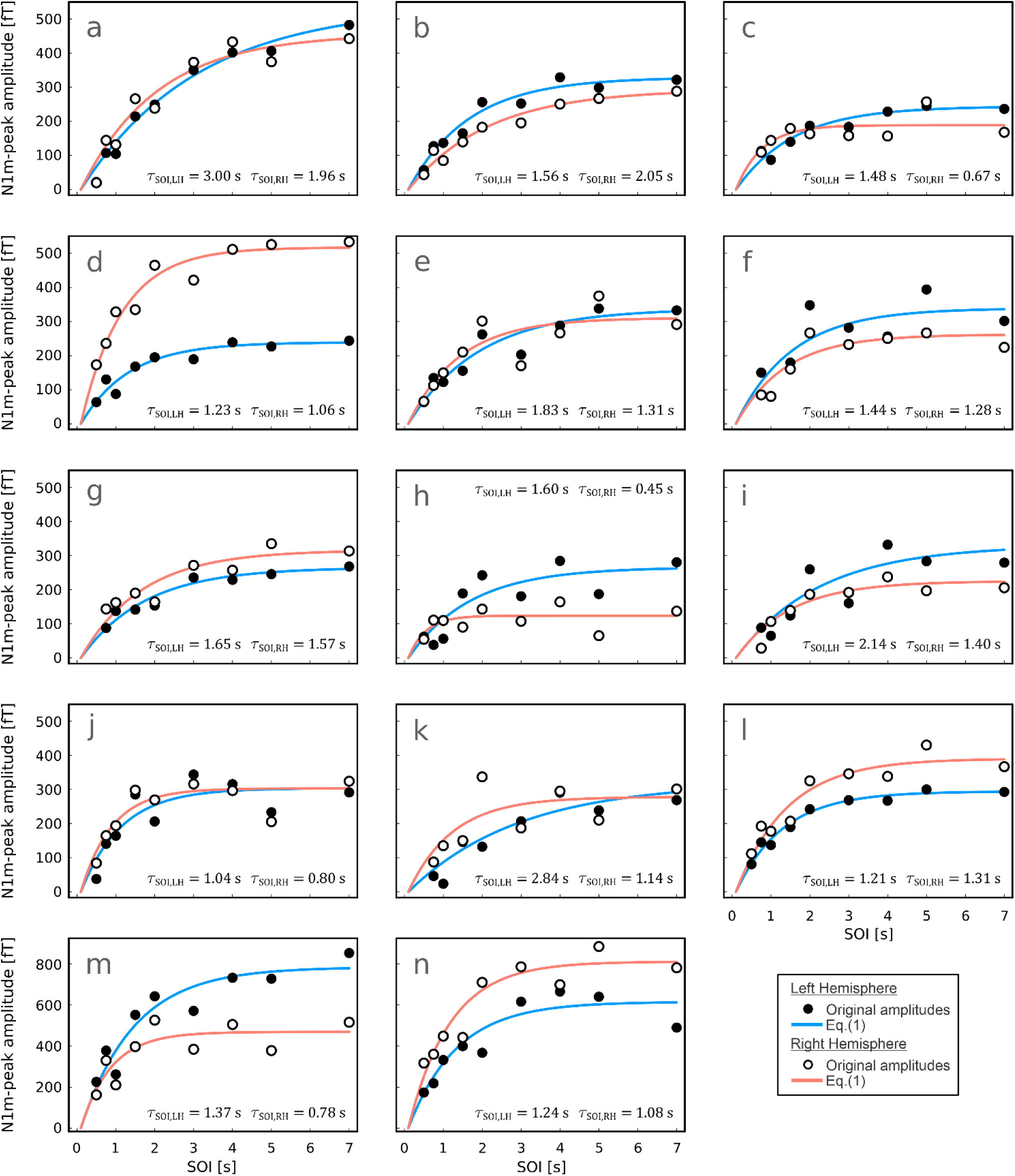
SOI-dependence of the N1m-peak amplitude for each subject. Each subplot shows the original N1m-peak amplitudes of the left (black solid symbols) and right (black open symbols) hemisphere. Blue and pink curves display Eq. (1) computed with the median τ_SOI_-and A-values. The respective hemisphere-specific values of τ_SOI_ are displayed in the figure panels. Note the different scale of the ordinate in the panels of subjects (m) and (n) compared to the other panels.

**Figure 3:**
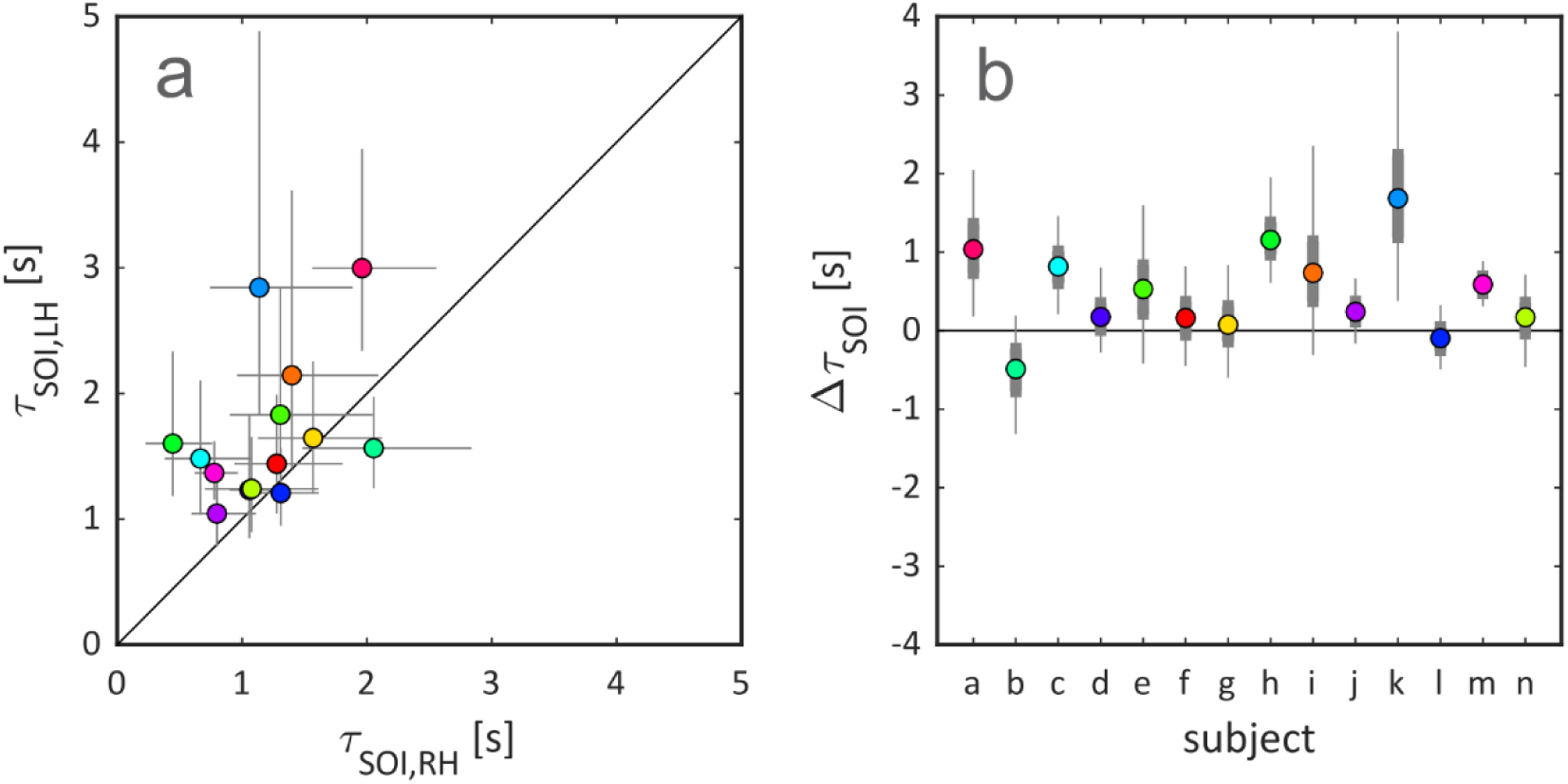
Subject-and hemisphere-specific results for the adaptation lifetime *τ*_SOI_. **a)** Comparison of *τ*_SOI_ between left and right hemisphere for all subjects of the cohort. Coloured symbols mark the median values for each subject, and the corresponding error bars oriented along each axis display the 95-% CIs. The diagonal line marks equal lifetimes for the two hemispheres. A Wilcoxon signed rank test revealed a hemisphere difference for *τ*_SOI_ with significantly larger values for the left hemisphere (*p* =0.004). **b)** Overview of Δ*τ*_SOI_-values across subjects. Coloured symbols (same colour code as in a) mark median values for each subject. The thicker grey error bars map out the interquartile range, and the thinner grey error bars the respective 95-% CIs. Median Δ*τ*_SOI_ exceeds zero in 12 out of 14 subjects. The subject labels at the abscissa are the same as in the panels of Fig. 2.

To explore the robustness of our finding that *τ*_SOI,LH_ > *τ*_SOI,RH_, we analysed the quantity Δ*τ*_SOI_ = *τ*_SOI,LH_ − *τ*_SOI,RH_. For each subject, we paired, randomly and without replacement, left and right hemisphere *τ*_SOI_-values obtained via the bootstrap approach and computed Δ*τ*_SOI_. Figure 3b depicts the resulting median values, interquartile ranges, and 95-% CIs of Δ*τ*_SOI_. The 95-% CIs include positive values across all 14 subjects. For subjects where the median Δ*τ*_SOI_-value, *m*(Δ*τ*_SOI_), exceeds zero (12 out of 14), even the lower quartile, *q*_1_(Δ*τ*_SOI_), exceeds zero in a majority of cases (8 out of 12, confirmed H1: where *m*(Δ*τ*_*SOI*_) > 0, *q*_1_(Δ*τ*_*SOI*_) ≠ 0, z = 2.27, p = 0.021). Out of all data points for Δ*τ*_SOI_, 78 % exceed zero. This confirms the robustness of our finding beyond the median values of Δ*τ*_*SOI*_.

The grand median of Δ*τ*_*SOI*_ across subjects, *M*(Δ*τ*_SOI_), is equal to 0.39 s, which amounts to 29 % of the grand median *τ*_SOI_-value across subjects and hemispheres, *M*(*τ*_SOI_) = 1.34 s. The lower quartile of the distribution, *Q*_1_(Δ*τ*_SOI_), is positive as well (0.16 s). This illustrates that the effect size of the hemispheric difference is not only large in terms of data ranks, but also translates into a substantial difference in terms of absolute values. Further, given the broad distributions of the bootstrap-based *τ*_SOI_-values derived for each subject, we also investigated the impact of outliers, defined as data points more than 1.5 interquartile ranges (IQRs) away from the lower or upper quartile (i.e. data point *x*_*i*_ is an outlier if *x*_*i*_ lies outside the interval [*q*_1_(*x*) − 1.5 · IQR(*x*), *q*_3_(*x*) + 1.5 · IQR(*x*)], where *x* denotes the subject-and hemisphere-specific distribution of *τ*_SOI_-values).

The removal of the outliers affected neither the grand median nor the test statistics for medians and lower quartiles of Δ*τ*_SOI_ substantially.

To summarise, Δ*τ*_SOI_ > 0 is a robust finding despite the large CIs for *τ*_SOI,LH_ and *τ*_SOI,RH_.

### 3.2 Saturation amplitude across subjects and hemispheres

Figure 4a shows the subject-specific median saturation amplitudes *A* for the two hemispheres (see Figure 2), and Figure 4b depicts the corresponding differences Δ*A* computed akin to Δ*τ*_SOI_, i.e. Δ*A* = *A*_LH_ − *A*_RH_. We observe that the boundaries of the range of median *A*-values for the left hemisphere (240 fT to 780 fT) are encompassed by those for the right hemisphere (120 fT to 810 fT). The same holds true for the ranges covered by the respective 95-% CIs, which extend from about 210 fT to 830 fT in the left and from 110 fT to 940 fT in the right hemisphere. This observation does not correspond to that for *τ*_SOI_ and the corresponding CIs, where the boundaries of the range of median values for the left hemisphere were larger than the respective counterparts for the right hemisphere (see Sect. 3.1). Further, the ratio between 95-%-CI ranges and respective medians for *A* was at least equal to 0.11, and at most equal to 0.54, with a grand median of 0.27.

**Figure 4:**
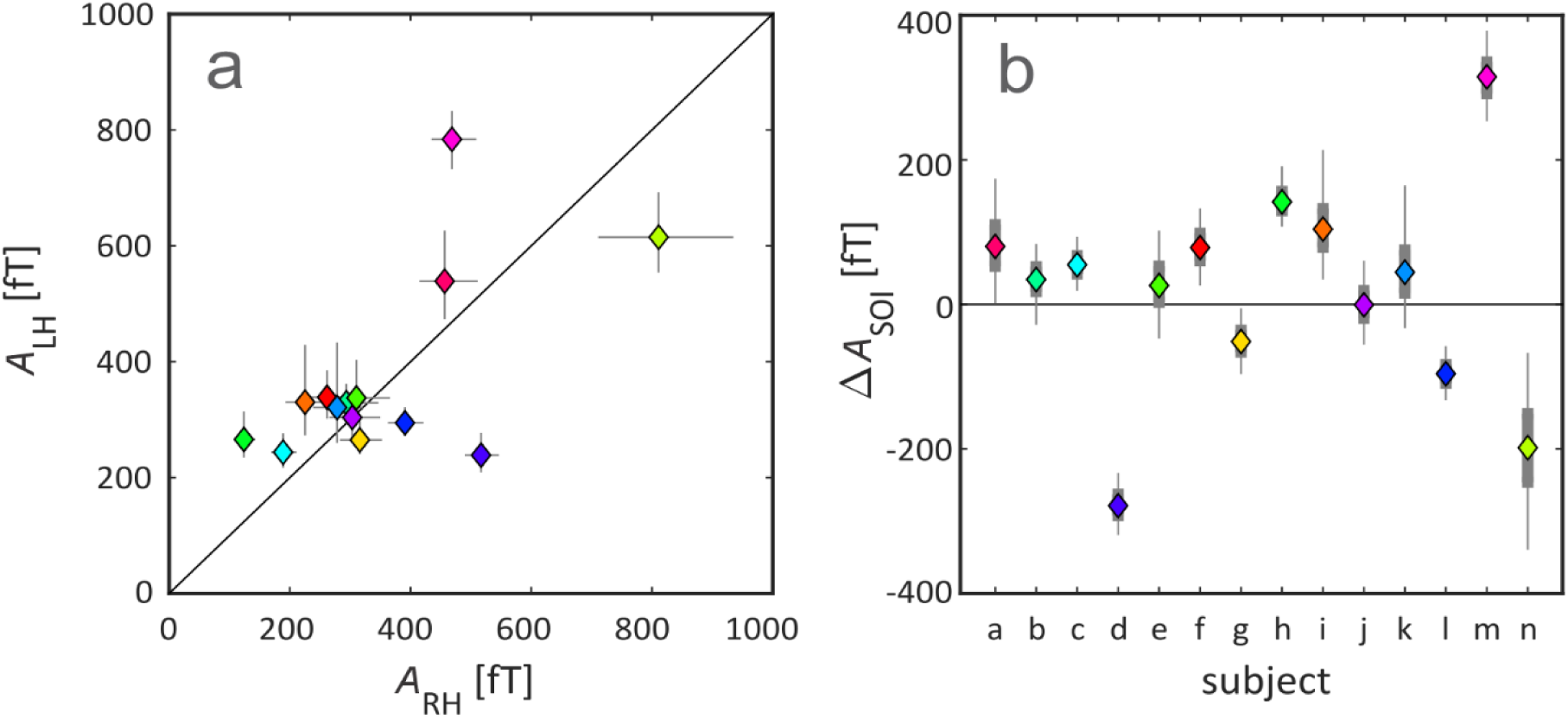
Subject-and hemisphere-specific results for the saturation amplitude *A*. **a)** Comparison of *A* between left and right hemisphere for all subjects. Coloured symbols mark the median values for each subject (same colour code as in Fig. 3a), and the corresponding error bars oriented along each axis display the 95-% CIs. A Wilcoxon signed rank test revealed no significant hemispheric difference for *A* (*p* = 0.426). The diagonal line marks equal amplitudes for the two hemispheres. **b)** Distribution of the bootstrap-based Δ*A*-values for each subject. Coloured symbols (same colour code as in a) mark the median for each subject. The thicker grey error bars map out the interquartile ranges, and the thinner grey error bars the respective 95-% CIs. The subject labels at the abscissa are the same as in the panels of Figures 2 and 3b.

Thus, overall, the relative size of the 95-%-CI ranges was much smaller for *A* than for *τ*_SOI_. A Wilcoxon signed rank test showed that there is no statistically significant difference between *A-* values of the left and right hemisphere (two-sided *p*-value = 0.426). Hence, the difference in the adaptation lifetime τ_SOI_ between left and right hemisphere (see Figure 3) is not accompanied by a difference in the saturation amplitude *A*.

### 3.3 Confirmation of results across different analysis conditions

We found that, when introducing intercept *t*_0_ as a third fitting parameter, the resulting median values stayed close to the fixed value *t*_0_ = 100 ms of the two-parameter fit. The grand median of *t*_0_ across both the baseline and non-baseline corrected analysis condition was equal to 137 ms (IQR: 100 ms to 328 ms) in the left and 255 ms (IQR: 100 ms to 351 ms) in the right hemisphere. Thus, deviations were small relative to the time scales observed for τ_SOI_. For the hypotheses that (1) there is a systematic difference between *τ*_SOI,LH_ and *τ*_SOI,RH_ (Δ*τ*_SOI_ ≠ 0) and (2) there is a systematic difference between *A*_LH_ and *A*_RH_ (Δ*A* ≠ 0), the *p*-values obtained across all four analysis conditions (with or without baseline correction and for a two-or three-parameter fit of Eq. (1)) are summarised in Table 1. As shown, the difference in τ_SOI_ across the two hemispheres is significant in all four conditions and the effect size remains large as well. Similarly, the lack of difference in *A*-values between left and right hemisphere is a robust finding, with differences remaining insignificant across all four conditions.

**Table 1:**
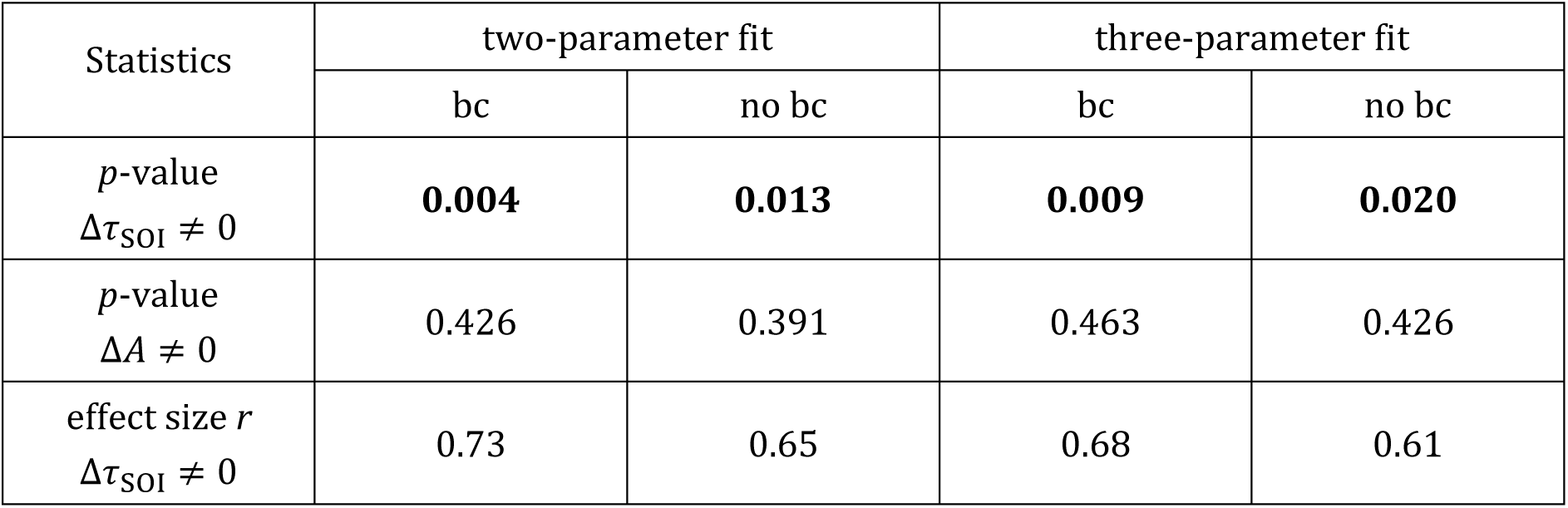
Summary of the *p*-values from the Wilcoxon signed rank tests for τ_SOI_ and *A* along with the corresponding effect size *r* of the result. Bold values indicate statistical significance. (bc = baseline correction)

| Statistics | two-parameter fit |  | three-parameter fit |  |
| --- | --- | --- | --- | --- |
|  | bc | no bc | bc | no bc |
| $p$-value<br>$\Delta\tau_{\text{SOI}} \neq 0$ | **0.004** | **0.013** | **0.009** | **0.020** |
| $p$-value<br>$\Delta A \neq 0$ | 0.426 | 0.391 | 0.463 | 0.426 |
| effect size $r$<br>$\Delta\tau_{\text{SOI}} \neq 0$ | 0.73 | 0.65 | 0.68 | 0.61 |

Taken together, these findings suggest that, within subjects, τ_SOI_ does not systematically increase with *A* across hemispheres.

### 3.4. Relationship between adaptation lifetime and saturation amplitude

To explore whether absolute values of τ_SOI_ and *A* co-vary across subjects, we investigated the relationship of these parameters across all subject-and hemisphere-specific median values. Figure 5a displays all median τ_SOI_-values as a function of the respective median *A*-values. The non-parametric Kendall rank correlation coefficient test confirmed that the correlation coefficient for this set of value pairs is very small and that the correlation is far from significance level (Kendall’s tau-b = 0.14, *p* = 0.317). This also applies at the hemisphere-specific level (left hemisphere: Kendall’s tau-b= 0.10, *p* = 0.667; right hemisphere: Kendall’s tau-b = 0.14, *p* = 0.518).

**Figure 5:**
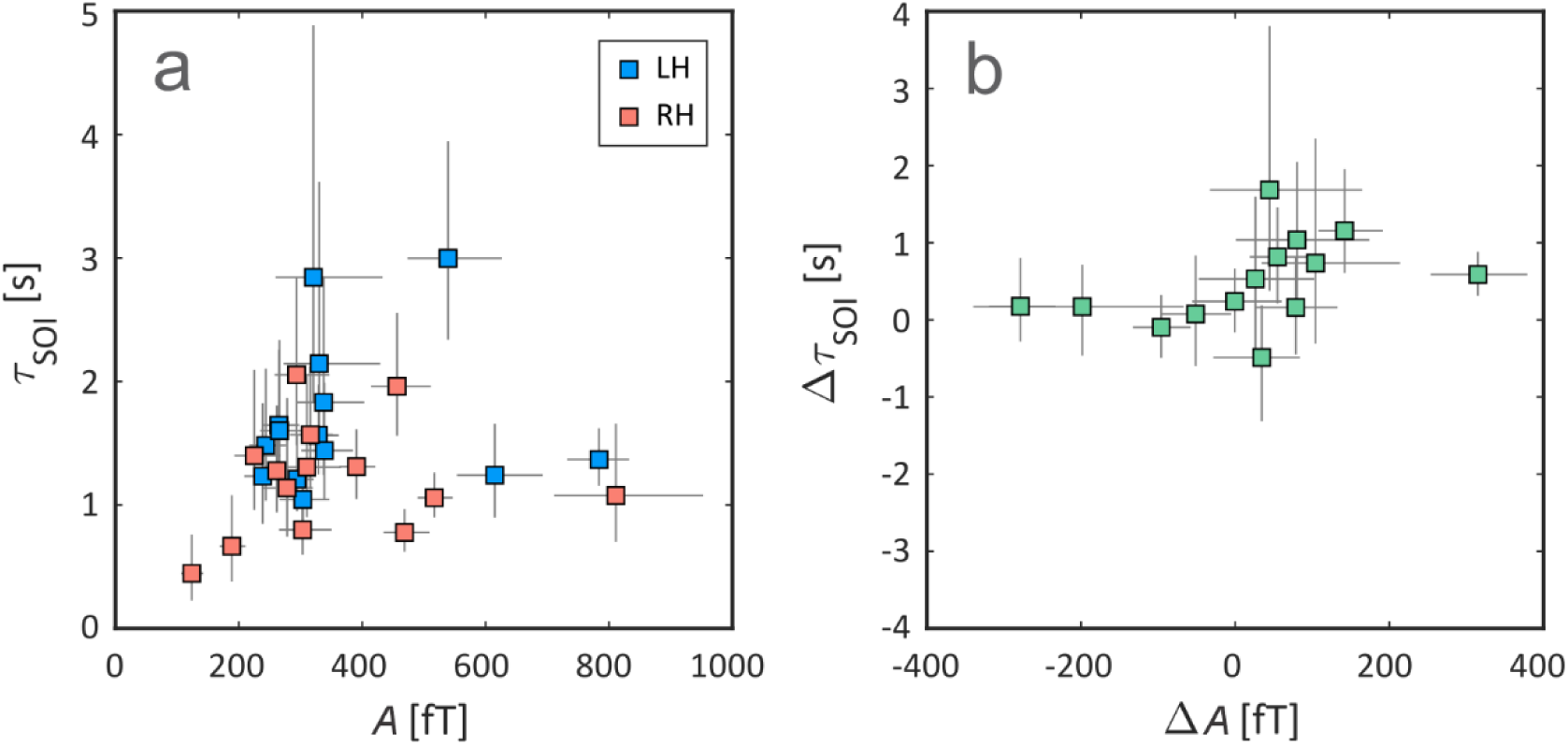
Relationship between adaptation lifetime and saturation amplitude. **a)** Adaptation lifetime *τ*_SOI_ as a function of saturation amplitude *A*. Blue and pink markers indicate the subject-specific median values for the left and right hemisphere, respectively. The error bars extend across the respective 95-% CI. There is no significant correlation between the median values for *τ*_SOI_ and *A* (*p* = 0.317). **b)** Difference between adaptation lifetimes, Δ*τ*_SOI_, as a function of the difference between saturation amplitudes, *ΔA*. The green markers map out subject-specific median Δ**τ**_SOI_-values, i.e. differences between left-and right-hemisphere adaptation lifetimes, as a function of subject-specific median Δ*A*-values, i.e. differences between left-and right-hemisphere saturation amplitudes of the N1m. The error bars extend across the respective 95-% CI. The covariance of the median Δ*τ*_SOI_-and *ΔA-*values is not statistically significant (*p* = 0.062).

To explore whether absolute values of τ_SOI_ and *A* co-vary across hemispheres, we investigated the relationship between Δ*τ*_SOI_ and Δ*A*. Figure 5b shows median Δ*τ*_SOI_ as a function of median Δ*A*, along with the respective 95-% CIs. Statistical testing showed no significant correlation between Δ*τ*_SOI_ and Δ*A* (Kendall’s tau-b = 0.38, p = 0.062).

We computed Kendall’s tau-coefficient for all 16 possible combinations of our analysis (τ_SOI_ vs. *A* for both hemispheres combined and for left and right hemisphere separately as well as Δ*τ*_SOI_ vs. Δ*A*, with and without baseline correction, two-and three-parameter fit of Eq. (1)). With two exceptions (1: τ_SOI_ vs. *A* in the right hemisphere, for a three-parameter fit *without baseline correction*, and 2: Δ*τ*_SOI_ vs Δ*A*, for a three-parameter fit *with baseline correction*), where *p* was minimally below the significance level, the *p*-values were always larger than 0.05.

To summarise, we confirm the hypothesis that τ_SOI_ and *A* do not co-vary, neither across subjects nor across hemispheres.

## 4 Discussion

### 4.1 Overview

Adaptation of neuronal responses is a basic feature of sensory processing in the brain, indexing the presence of sensory memory. Here, we explored by means of MEG whether the adaptation lifetime *τ*_SOI_ shows asymmetries across the left and right AC. We used binaural stimulation and the regular-SOI paradigm to determine the adaptation lifetime *τ*_SOI_ expressed in terms of the N1m response of the auditory ERF and developed a bootstrap-based analysis pipeline for subject-specific statistical inference. Median subject-specific *τ*_SOI_-values ranged from about 1.0 s to 3.0 s in the left hemisphere, and from 0.4 s to 2.0 s in the right. The grand median across subjects and hemispheres, *M*(*τ*_SOI_), was 1.34 s. Our data showed that, at the single-subject level, there is a statistically significant difference, Δ*τ*_SOI_, between *τ*_SOI_ in the left and right hemisphere. The grand median of this difference, *M*(Δ*τ*_SOI_), was 0.39 s, i.e. 29 % of *M*(*τ*_SOI_). In contrast, there was no significant hemispheric difference in the saturation of the N1m-peak amplitude *A*, and no significant correlation between *τ*_SOI_ and *A*. Therefore, the hypothesis that *τ*_SOI_ varies as a function of *A* was rejected. To our knowledge, this is the first demonstration of a hemispheric difference in the adaptation lifetime of the human AC.

### 4.2 Comparison with results of previous studies on adaptation lifetime

In ERF and ERP studies, the determination of adaptation lifetime is commonly based on the SOI-dependence of the peak amplitude of the N1m/N1. Hence, to study the hemispheric asymmetry of *τ*_SOI_, one must take in account the hemispheric differences in the N1m/N1 response that depend on whether the stimulus is presented monaurally or binaurally. The N1m/N1-response was reported to be larger in the hemisphere contralateral to the stimulated ear in studies with monaural (see, for example, Hine & Debener, 2007; McEvoy et al., 1997; Pantev et al., 1998; Reite et al., 1981; Ross et al., 2005; Salmelin et al., 1999; Woldorff et al., 1999) or alternating binaural stimulation (Mäkelä et al., 1993). Specifically, Hine & Debener (2007) reported that EEG-recorded activity was significantly larger in the right than in the left hemisphere when the stimuli were presented to the respective contralateral ear (see also Kanno et al., 1996). This general right-hemispheric preponderance is commonly attributed to the difference in cortical folding between the two hemispheres, which results in larger signal cancelation in the left hemisphere (Shaw et al., 2013). However, with binaural stimulation, Ross et al. (2005) found no pronounced hemispheric difference for the amplitude of the grand mean source waveforms of the ERF, including N1m.

While previous studies have reported separate adaptation lifetimes for the left and right AC, these experiments usually used monaural stimulation and/or tended to be of low power (c.f. literature cited in the Introduction). The issue with monaural stimulation is that the contralaterally stronger responses could confound any hemispheric differences in *τ*_SOI_. In consequence, no robust conclusions regarding hemispheric asymmetry of adaptation could be drawn. In the following paragraphs, we discuss our results in the context of previous studies with binaural stimulation reported in the literature.

Mäkelä et al. (1993) recorded ERFs from nine subjects where tones (1-kHz audio-frequency, 50-ms duration, 80-85-dB hearing level) were delivered *alternately* to the two ears at five different interstimulus intervals of 1, 2, 4, 8, and 16 s. Setting *t*_0_ = 0 s in Eq. (1), i.e. using a two-parameter approach with the intercept of the fit at the origin, the authors report that the average dependence of the N1m-peak response on the interstimulus interval was similar over both hemispheres and for both contralateral and ipsilateral stimuli. They report *τ*_SOI_-values of 0.9 s and 1.5 s for the contra-and ipsilateral N100m in the right hemisphere and 1.2 s and 1.5 s for the corresponding values in the left hemisphere, which are in a similar range as those reported in this study.

Cheng & Lin (2012) studied the effects of aging on the lifetime of adaptation in humans. The authors recorded N1m responses in young, middle-aged, and elderly subjects (n = 15 for each group), who were exposed to trains of five successive tones (800-Hz audio-frequency, 25-ms duration, 65-70 dB intensity above hearing threshold). In separate sessions, five within-train SOIs of 0.5 s, 1 s, 2 s, 4 s, and 8 s were presented; the inter-train interval was 10 s. The adaptation time constant was determined from the amplitude ratio between the N1m-peak responses to the fifth (last) and the first tone (S5/S1 in the author’s notation) of the train according to Eq. (1). The authors report a decrease of mean *τ*_SOI_-values with increasing age, with *τ*_SOI_ = (1.06 ± 0.26) s for the elderly, *τ*_SOI_ = (1.70 ± 0.25) s for the middle-aged and the *τ*_SOI_ = (2.77 ± 0.25) s for the young subjects; however, no statistically significant differences between the left and the right hemisphere were found in any group. The reason for the discrepancy between the absence of a hemispheric difference of *τ*_SOI_ in Cheng & Lin’s results compared to our findings might be traced back to differences in the experimental design and the related analysis. Essential aspects could be that Cheng & Lin used only five SOIs in their study, which clearly affects the quality of the exponential fit of the SOI-dependence of the N1m-peak amplitude. Further, they recorded only 20-25 repetitions for each stimulus, which often resulted in low SNRs. This may also be the cause why the amplitude ratio S5/S1 is greater than 1 in many cases, especially for large SOIs.

Ioannides et al. (2003) used MEG to investigate the interhemispheric asymmetry in adaptation lifetime and its dependence on handedness and gender. Subjects listened to pure tones (1-kHz audio-frequency, 200-ms duration, 60-dB nHL) that were presented binaurally in four separate runs. Each run contained 30 repetitions of a block of stimuli, and in each block, nine tones were spaced in a fixed sequence using six SOIs (1 s, 1.5 s, 2 s, 3 s, 5 s, and 8 s). The sequence of the tones differed between the four runs. Using this type of *random-SOI* paradigm together with a more complex analysis that takes into account the entire N1m waveform, *τ*_SOI_ estimated at the peak of the N1m was found to be very similar in the left (1.6 s) and right (1.4 s) hemisphere. In contrast, adaptation lifetime estimated on the descending phase of N1m was much longer in the left hemisphere than in the right. It is important to note that in about half of the subjects SOI = 8 s or SOIs = 5 s and 8 s had to be excluded from the analysis, thus reducing the number of SOIs available for the exponential fit to 5 or under.

In conclusion, the absence of a significant hemispheric difference in adaptation lifetime previously reported in the literature may have at least partly resulted from the boundary conditions of the experimental setup and/or from the selected analysis approach. The results presented in the current study are based on the regular-SOI, not the random-SOI paradigm. They suggest that a hemispheric difference in adaptation lifetime exists in the context of this paradigm.

This finding can be considered particularly reliable for several reasons: 1) the number of stimulus repetitions (≥100), SOIs (9), and subjects (14) was comparatively large, 2) the hemispheric difference was observed for both two-as well as three-parameter exponential fits of Eq. (1), and 3) the bootstrap pipeline confirmed the robustness of this result.

### 4.3 Methodological challenges in the determination of the adaption lifetime

As expected, the noise sensitivity of the MEG signal proved to be a major challenge in the investigation. In a regular-SOI paradigm, the signal-to-noise ratio of ERFs can be very poor unless hundreds of trials are averaged. However, a substantial increase in the number of trials for more efficient noise suppression is only feasible with short SOIs. With long SOIs of several seconds, the use of even 100 stimulus repetitions leads to excessive measurement durations that are unacceptable in terms of subject endurance and can lead to data distortions due to fatigue. Against this background, the bootstrap technique opens up an auspicious solution to this quandary, as the approach generates additional surrogate ERFs based on existing MEG data. Consequently, the number of bootstrapped ERFs per SOI (999 in our analysis) can exceed the number of original ERFs many times over. This yields information about the stability of the results and allows for statistical inferences about the adaptation lifetime, all while requiring neither a normal distribution nor homoscedasticity of the original data.

Traditionally, the adaptation lifetime is derived by fitting a single exponential function to the SOI-dependence of the peak amplitude of the N1m, as implemented by, for example, Lu et al. (1992). However, single exponential functions are very rare in nature, and this may also apply to the adaptation process in AC as discussed here. Does a single exponential function, and thus a single time constant, sufficiently explain the process of adaptation recovery? Hajizadeh et al. (2022) raised this question in their computational modelling study addressing ERF generation and adaptation. The authors concluded that special care must be taken with regard to the range of sampling points when a single exponential function is fitted to the data. Alternatively, at the cost of increasing the number of fitting parameters, the use of other fitting functions such as the stretched exponential Kohlrausch function could be an option (Lukichev, 2019). Regardless of which function is used, the purpose of such a function is to provide an estimate of the speed with which neuronal responses recover from adaptation. While these estimates are necessarily crude—due also to the noise in the data—they are required for making comparisons of adaptation lifetime across participants, hemispheres, and experimental conditions. How any estimating function can be linked to the underlying neurophysiological processes remains an open question.

An additional complication in deriving adaptation lifetime arises when entering the realm of very short SOIs. The change in response behaviour in this SOI-range is particularly evident in the random-SOI paradigm. In this paradigm, the SOI between successive stimuli is randomly assigned and single-trial responses are sorted and averaged according to the immediately preceding SOI. In this case, Zacharias et al. (2012) observed an N1m-*facilitation* at SOIs below ∼0.5 s. The peak amplitude *decreases* as a function of SOI up to about the 0.5-s mark and only starts to increase (i.e. to recover from adaptation) for SOIs larger than ∼0.5 s (see also, for example, Budd and Michie 1994; Loveless et al. 1996; Sable et al. 2004; Wang et al. 2008). This indicates that, at short SOIs, the N1m is sensitive to the stimulation history extending further back than to the previous stimulus. In this light, the omission of the 250-ms SOI in our analyses of data recorded with the regular-SOI paradigm allowed us to bypass possible ambiguities in the calculation of *τ*_SOI_.

We derived the hemisphere-specific adaptation lifetimes using measurements from a single channel per hemisphere, that is, from the channel displaying the largest N1m amplitude. We purposely dispensed with source modelling, the iterative search for a solution of the inverse problem in terms of point-like equivalent current dipoles (ECDs). In classical source modelling, one or two ECDs per hemisphere are used to describe event-related AC activity. Although already a single ECD per hemisphere can often be considered an adequate solution, it is unable to account for observations such as N1m responses with double peaks, which calls for a two-ECD model. However, in our study, the application of a model that comprises two ECDs per hemisphere would be far from straightforward. On the one hand, the source-modelling algorithm in this case must be able to separate two sources that are closely linked both spatially and temporally. This is by no means an easy task. On the other hand, by the time the N1m emerges, activation has spread to large extents of AC, including belt and parabelt areas. This is evident in experimental findings (for a review, see May & Tiitinen, 2010) and is also indicated by computational modelling of AC (Hajizadeh et al., 2019). This means that two dipoles is just as unrealistic a model of N1m generation as a single dipole. We therefore do not expect ECD source modelling to lead to more meaningful findings than the analysis of the measured magnetic fields carried out here.

### 4.4 Lessons from computational modelling on adaptation lifetime variations

The difference of adaptation lifetime across the hemispheres could be due to several reasons. May and colleagues (2015) simulated a single AC in a computational model which encapsulates the large-scale organisation of AC into core, belt, and parabelt fields. They found that the adaptation of the N1m, as well as selective responding to complex tone sequences, can be explained as resulting from short-term synaptic depression (STSD). Further, the kinetics of this depression determined the adaptation lifetime of the N1m. The findings reported in this work could therefore arise from slower STSD decay in the left hemisphere than in the right.

Hajizadeh et al. (2022), working with a similar model as May and colleagues, found that the SOI-specific response behaviour of the AC also varies as a function of cortical folding and of network connectivity. Their model allowed the description of AC dynamics in terms of overlapping normal modes, each with its own frequency and damping, and where each normal mode is spatially distributed over the entire system (see also Hajizadeh et al., 2019). The ERF signal as a whole is a linear, weighted sum of these normal modes. Two important criteria that arise from the model equations dictate the composition of the ERF: the input efficiency and the MEG efficiency. The input efficiency determines to which extent input from subcortical regions activates each normal mode, and the MEG efficiency determines to which extent the MEG device detects this activation. One parameter that modulates MEG efficiency is cortical folding. Anatomical studies have shown that folding of the AC is hemisphere-specific (Heschl, 1878; Morosan et al., 2001; Rademacher et al., 2001; v. Economo & Horn, 1930). In their simulations, Hajizadeh et al. (2021) showed that such differences in cortical folding can lead to changes in ERF waveforms, in particular in terms of N1m-peak amplitudes. Thus, hemisphere-specific cortical folding could cause differences in adaptation lifetimes, even if STSD kinetics are identical across hemispheres. Moreover, MEG efficiency and input efficiency, as well as the normal modes themselves, vary as a function of network connectivity. Thus, even if STSD dynamics *and* cortical folding are identical across hemispheres, differences in adaptation lifetime could arise via differences in connection pattern and strengths. Hajizadeh et al. (2022) found that by changing which AC fields were connected to each other, i.e. by changing the connection *pattern*, the adaptation lifetime *τ*_SOI_ estimated from the N1m peaks was modified from 2.3 s to 2.7 s. The effect was thus of the same magnitude as observed in the current study.

Further, Hajizadeh et al. (2022) found that, in their model, the factors that govern the superposition of network activity into ERFs—the MEG efficiency and the input efficiency introduced above—are SOI-specific. This could mean that the jitter of the SOI-specific N1m-peak amplitudes around the fitted curve, as depicted in Figure 2, is not just due to noise but also reflects an actual SOI-specific effect. Previous studies have shown that subject-specific ERF waveforms are very stable across days, weeks, and even years (see, e.g., Ahonen et al., 2016; Atcherson et al., 2006; Sandman & Patterson, 2000; Segalowitz & Barnes, 1993). Thus, one way of verifying this hypothesis would be to repeat the experiment described in this work with the same subjects and then compare the ‘jitter pattern’. If the hypothesis were to be confirmed, no monotonically rising fitting function could fully reflect the ACs’ recovery from adaptation as a function of SOI.

Hajizadeh et al.’s finding that N1m adaptation depends on AC connectivity is supported by simulation results reported by Tomana et al. (2023). Using a simplified version of the model by May and colleagues and an optimisation approach based on an evolutionary algorithm, the authors showed that, even when preserving the connection pattern, changes to model parameters reflecting connection *strengths* lead to changes in N1m adaptation. The optimisation of the values for these parameters strongly increased the overlap between simulated and experimental auditory ERFs across a 250-ms time window. Adaptation lifetime in the best-performing version of the initial model was characterised by a *τ*_SOI_ of 1.8 s whereas in the optimised model it was characterised by a *τ*_SOI_ of 2.0 s. Thus, it is possible that the hemispheric differences in adaptation that we report here are at least partly explained by the left AC being differently organised than its right counterpart in terms of connections patterns and connections strengths.

In summary, previous computational modelling suggests that the observed difference in adaptation lifetime across the cortical hemispheres can be explained by at least three factors: 1) hemisphere-specific time scales for the underlying synaptic plasticity that gives rise to the adaptation phenomenon, 2) hemisphere-specific cortical folding that affects the relationship between neural activation patterns and extracranial magnetic fields, or 3) hemisphere-specific AC connectivity that modulates the SOI-specific response behaviour of the network. Note that the above explanations are not mutually exclusive, and that the contribution of each to observed adaptation lifetime remains to be explored.

### 4.5 Implications of hemisphere-specific adaptation lifetimes

Auditory stimulation, especially in the form of speech, can be characterised by a range of different features that evolve along diverging time scales (see, e.g., Rosen et al., 1997). Information regarding individual phonemes and short syllables, for example, is encompassed by very short time windows of about 20 ms to 40 ms. In contrast, information regarding intonation is contained in longer time windows with a duration of about 150 ms to 250 ms. The processing of fast transitions at time scales characteristic for phonemes and short syllables was found to cause cortical activity lateralised to the left hemisphere, whereas tasks involving information evolving on a longer scale, such as intonation and prosody, lead to stronger activation of the right hemisphere (Arnal et al., 2015; Poeppel, 2003). Poeppel, Arnal, and colleagues suggest that this asymmetry arises due to an ‘asymmetric sampling in time’ across the two hemispheres.

In contrast to phonemes and intonation, the adaptation phenomenon studied in this work is based on the auditory processing of pure tones, and characterised by a much larger time scale of the order of seconds. Nevertheless, at this different time scale, our results potentially add to the overall picture of asymmetries in the temporal processing of auditory stimulation. Lu et al. (1992) showed that subject-specific adaptation lifetimes, also based on pure-tone stimulation and quantified in the same manner as in our work, correlate with behaviourally observed subject-specific lifetimes of an auditory sensory memory trace. Thus, τ_SOI_ could be viewed as a measure for the size of the time window available for the adaptation-based integration of auditory information. Although the time windows associated with the left and right auditory cortex largely overlap, we also identified a significant difference: the grand median difference across the hemispheres was almost 400 ms, with longer time windows in the left hemisphere. This difference might have a functional relevance in the context of the lateralisation of temporal binding.

In this context, our results suggest that a lateralisation of information processing, i.e. a form of ‘asymmetric sampling in time’, might also be observed for different stimulus features evolving at a time scale of seconds rather than milliseconds. This in turn raises the question whether the asymmetry we observed is retained or boosted if the stimulation is task-relevant and therefore in the focus of selective attention. Finally, in the context of the work by Lu and colleagues (1992), it also needs to be established whether adaptation in the left or the right auditory cortex better predicts sensory memory lifetime as estimated with behavioural methods.

## Acknowledgements

We would like to thank Artur Matysiak for his support in taking the MEG measurements and in developing the bootstrap-based analysis. This research was supported by the Deutsche Forschungsgemeinschaft under project no. K01713/12-1.

## References

Ahonen, L., Huotilainen, M., & Brattico, E. (2016). Within-and between-session replicability of cognitive brain processes: An MEG study with an N-back task. Physiology & Behavior, 158, 43–53. 10.1016/J.PHYSBEH.2016.02.006

Arnal, L. H., Poeppel, D., & Giraud, A. (2015). Temporal coding in the auditory cortex. In Handbook of Clinical Neurology (Vol. 129, pp. 85–98). Elsevier. 10.1016/B978-0-444-62630-1.00005-6

Atcherson, S. R., Gould, H. J., Pousson, M. A., & Prout, T. M. (2006). Long-Term Stability of N1 Sources Using Low-Resolution Electromagnetic Tomography. Brain Topography, 19(1), 11–20. 10.1007/s10548-006-0008-8

Bezanson, J., Edelman, A., Karpinski, S., & Shah, V. B. (2017). Julia: A Fresh Approach to Numerical Computing. SIAM Review, 59(1), 65–98. 10.1137/141000671

Brechmann, A., & Scheich, H. (2005). Hemispheric shifts of sound representation in auditory cortex with conceptual listening. *Cerebral Cortex (New York*, N.Y.: 1991), *15*(5), 578–587. 10.1093/cercor/bhh159

Broca, M. P. (1861). Remarks on the seat of the faculty of articulated language, following an observation of aphemia (loss of speech). Bulletin de la Société Anatomique, 6, 330–357.

Budd, T. W., & Michie, P. T. (1994). Facilitation of the N1 peak of the auditory ERP at short stimulus intervals. NeuroReport, 5(18), 2513–2516.

Butler, R. A. (1968). Effect of changes in stimulus frequency and intensity on habituation of the human vertex potential. The Journal of the Acoustical Society of America, 44(4), 945–950. 10.1121/1.1911233

Cha, K., Zatorre, R. J., & Schönwiesner, M. (2016). Frequency Selectivity of Voxel-by-Voxel Functional Connectivity in Human Auditory Cortex. Cerebral Cortex, 26(1), 211–224. 10.1093/cercor/bhu193

Cheng, C.-H., & Lin, Y.-Y. (2012). The effects of aging on lifetime of auditory sensory memory in humans. Biological Psychology, 89(2), 306–312. 10.1016/j.biopsycho.2011.11.003

Dalebout, S. D., & Robey, R. R. (1997). Comparison of the intersubject and intrasubject variability of exogenous and endogenous auditory evoked potentials. Journal of the American Academy of Audiology, 8(5), 342–354.

Efron, B. (1979). Bootstrap Methods: Another Look at the Jackknife. The Annals of Statistics, 7(1), 1–26. 10.1214/AOS/1176344552

Friston, K. (2005). A theory of cortical responses. Philosophical Transactions of the Royal Society B: Biological Sciences, 360(1456), 815–836. 10.1098/rstb.2005.1622

Fritz, C. O., Morris, P. E., & Richler, J. J. (2012). Effect size estimates: Current use, calculations, and interpretation. Journal of Experimental Psychology: General, 141(1), 2–18. 10.1037/a0024338

Hajizadeh, A., Matysiak, A., Brechmann, A., König, R., & May, P. J. C. (2021). Why do humans have unique auditory event-related fields? Evidence from computational modeling and MEG experiments. Psychophysiology, 58(4), e13769. 10.1111/psyp.13769

Hajizadeh, A., Matysiak, A., May, P. J. C., & König, R. (2019). Explaining event-related fields by a mechanistic model encapsulating the anatomical structure of auditory cortex. Biological Cybernetics, 113(3), 1–25. 10.1007/s00422-019-00795-9

Hajizadeh, A., Matysiak, A., Wolfrum, M., May, P. J. C., & König, R. (2022). Auditory cortex modelled as a dynamical network of oscillators: Understanding event-related fields and their adaptation. Biological Cybernetics, 116(4), 475–499. 10.1007/ S00422-022-00936-7

Heschl, R.-L. (1878). Über die vordere quere Schläfenwindung des menschlichen Gehirns. Braumüller., Wien

Hine, J., & Debener, S. (2007). Late auditory evoked potentials asymmetry revisited. Clinical Neurophysiology, 118(6), 1274–1285. 10.1016/j.clinph.2007.03.012

Ioannides, A. A., Popescu, M., Otsuka, A., Bezerianos, A., & Liu, L. (2003). Magnetoencephalographic evidence of the interhemispheric asymmetry in echoic memory lifetime and its dependence on handedness and gender. NeuroImage, 19(3), 1061–1075. 10.1016/s1053-8119(03)00175-7

Kanno, A., Nakasato, N., Fujita, S., Seki, K., Kawamura, T., Ohtomo, S., Fujiwara, S., & Yoshimoto, T. (1996). Right hemispheric dominance in the auditory evoked magnetic fields for pure-tone stimuli. Electroencephalography and Clinical Neurophysiology. Supplement, 47, 129– 132.

Leech, R., Holt, L. L., Devlin, J. T., & Dick, F. (2009). Expertise with Artificial Nonspeech Sounds Recruits Speech-Sensitive Cortical Regions. Journal of Neuroscience, 29(16), 5234–5239. 10.1523/JNEUROSCI.5758-08.2009

Loveless, N., Levänen, S., Jousmäki, V., Sams, M., & Hari, R. (1996). Temporal integration in auditory sensory memory: Neuromagnetic evidence. Electroencephalography and Clinical Neurophysiology, 100(3), 220–228. 10.1016/0168-5597(95)00271-5

Lü, Z. L., Williamson, S. J., & Kaufman, L. (1992). Human auditory primary and association cortex have differing lifetimes for activation traces. Brain Research, 572(1–2), 236–241. 10.1016/0006-8993(92)90475-O

Lu, Z., Williamson, S., & Kaufman, L. (1992). Behavioral lifetime of human auditory sensory memory predicted by physiological measures. Science, 258(5088), 1668–1670. 10.1126/science.1455246

Lukichev, A. (2019). Physical meaning of the stretched exponential Kohlrausch function. Physics Letters A, 383(24), 2983–2987. 10.1016/j.physleta.2019.06.029

Mäkelä, J. P., Ahonen, A., Hämäläinen, M., Hari, R., Llmoniemi, R., Kajola, M., Knuutila, J., Lounasmaa, O. V., McEvoy, L., Salmelin, R., Salonen, O., Sams, M., Simola, J., Tesche, C., & Vasama, J.-P. (1993). Functional differences between auditory cortices of the two hemispheres revealed by whole-head neuromagnetic recordings. Human Brain Mapping, 1(1), 48–56. 10.1002/hbm.460010106

May, P. J. C. (2021). The Adaptation Model Offers a Challenge for the Predictive Coding Account of Mismatch Negativity. Frontiers in Human Neuroscience, 15, 721574. 10.3389/fnhum.2021.721574

May, P. J. C., & Tiitinen, H. (2010). Mismatch negativity (MMN), the deviance-elicited auditory deflection, explained. Psychophysiology, 47(1), 66–122. 10.1111/j.1469-8986.2009.00856.x

May, P. J. C., Westö, J., & Tiitinen, H. (2015). Computational modelling suggests that temporal integration results from synaptic adaptation in auditory cortex. The European Journal of Neuroscience, 41(5), 615–630. 10.1111/ejn.12820

McEvoy, L., Levänen, S., & Loveless, N. (1997). Temporal characteristics of auditory sensory memory: Neuromagnetic evidence. Psychophysiology, 34(3), 308–316. 10.1111/J.1469-8986.1997.TB02401.X

Michalewski, H. J., Prasher, D. K., & Starr, A. (1986). Latency variability and temporal interrelationships of the auditory event-related potentials (N1, P2, N2, and P3) in normal subjects. Electroencephalography and Clinical Neurophysiology/Evoked Potentials Section, 65(1), 59–71. 10.1016/0168-5597(86)90037-7

Morosan, P., Rademacher, J., Schleicher, A., Amunts, K., Schormann, T., & Zilles, K. (2001). Human primary auditory cortex: Cytoarchitectonic subdivisions and mapping into a spatial reference system. NeuroImage, 13(4), 684–701. 10.1006/nimg.2000.0715

Näätänen, R., Gaillard, A. W., & Mäntysalo, S. (1978). Early selective-attention effect on evoked potential reinterpreted. Acta Psychologica, 42(4), 313–329. 10.1016/ 0001-6918(78)90006-9

Näätänen, R., & Picton, T. (1987). The N1 Wave of the Human Electric and Magnetic Response to Sound: A Review and an Analysis of the Component Structure. Psychophysiology, 24(4), 375–425. 10.1111/j.1469-8986.1987.tb00311.x

Näätänen, R., Schröger, E., Karakas, S., Tervaniemi, M., & Paavilainen, P. (1993). Development of a memory trace for a complex sound in the human brain. NeuroReport, 4(5), 503–506.

Ocklenburg, S., & Güntürkün, O. (2024). Language and the left hemisphere. In The Lateralized Brain (pp. 129–165). Elsevier. 10.1016/B978-0-323-99737-9.00010-0

Pantev, C., Ross, B., Berg, P., Elbert, T., & Rockstroh, B. (1998). Study of the human auditory cortices using a whole-head magnetometer: Left vs. right hemisphere and ipsilateral vs. contralateral stimulation. Audiology & Neuro-Otology, 3(2–3), 183–190. 10.1159/000013789

Parras, G. G., Nieto-Diego, J., Carbajal, G. V., Valdés-Baizabal, C., Escera, C., & Malmierca, M. S. (2017). Neurons along the auditory pathway exhibit a hierarchical organization of prediction error. Nature Communications, 8(1), 2148. 10.1038/ s41467-017-02038-6

Pérez-González, D., & Malmierca, M. S. (2014). Adaptation in the auditory system: An overview. Frontiers in Integrative Neuroscience, 8, 19. 10.3389/ fnint.2014.00019

Poeppel, D. (2003). The analysis of speech in different temporal integration windows: Cerebral lateralization as “asymmetric sampling in time.” Speech Communication, 41(1), 245–255. 10.1016/S0167-6393(02)00107-3

Rademacher, J., Morosan, P., Schleicher, A., Freund, H.-J., & Zilles, K. (2001). Human primary auditory cortex in women and men. NeuroReport, 12(8), 1561–1565.

Rao, R. P. N., & Ballard, D. H. (1999). Predictive coding in the visual cortex: A functional interpretation of some extra-classical receptive-field effects. Nature Neuroscience, 2(1), 79–87. 10.1038/4580

Reite, M., Zimmerman, J. T., & Zimmerman, J. E. (1981). Magnetic auditory evoked fields: Interhemispheric asymmetry. Electroencephalography and Clinical Neurophysiology, 51(4), 388–392. 10.1016/0013-4694(81)90102-4

Rojas, D. C., Teale, P., Sheeder, J., & Reite, M. (1999). Sex differences in the refractory period of the 100 ms auditory evoked magnetic field. Neuroreport, 10(16), 3321–3325. 10.1097/00001756-199911080-00013

Rosburg, T., Zimmerer, K., & Huonker, R. (2010). Short-term habituation of auditory evoked potential and neuromagnetic field components in dependence of the interstimulus interval. Experimental Brain Research, 205(4), 559–570. 10.1007/ S00221-010-2391-3

Rosen, S., Carlyon, R. P., Darwin, C. J., & Russell, I. J. (1997). Temporal information in speech: Acoustic, auditory and linguistic aspects. Philosophical Transactions of the Royal Society of London. Series B: Biological Sciences, 336(1278), 367–373. 10.1098/ rstb.1992.0070

Ross, B., Herdman, A. T., & Pantev, C. (2005). Right hemispheric laterality of human 40 Hz auditory steady-state responses. Cerebral Cortex, 15(12), 2029–2039. 10.1093/ cercor/bhi078

Ruthig, P., & Schönwiesner, M. (2022). Common principles in the lateralization of auditory cortex structure and function for vocal communication in primates and rodents. European Journal of Neuroscience, 55(3), 827–845. 10.1111/ejn.15590

Sable, J. J., Low, K. A., Maclin, E. L., Fabiani, M., & Gratton, G. (2004). Latent inhibition mediates N1 attenuation to repeating sounds. Psychophysiology, 41(4), 636–642. 10.1111/j.1469-8986.2004.00192.x

Salmelin, R., Schnitzler, A., Parkkonen, L., Biermann, K., Helenius, P., Kiviniemi, K., Kuukka, K., Schmitz, F., & Freund, H. (1999). Native language, gender, and functional organization of the auditory cortex. Proceedings of the National Academy of Sciences of the United States of America, 96(18), 10460–10465. 10.1073/pnas.96.18.10460

Sandman, C. A., & Patterson, J. V. (2000). The auditory event-related potential is a stable and reliable measure in elderly subjects over a 3 year period. Clinical Neurophysiology, 111(8), 1427–1437. 10.1016/S1388-2457(00)00320-5

Schönwiesner, M., Rübsamen, R., & Von Cramon, D. Y. (2005). Hemispheric asymmetry for spectral and temporal processing in the human antero-lateral auditory belt cortex. European Journal of Neuroscience, 22(6), 1521–1528. 10.1111/j.1460-9568.2005.04315.x

Segalowitz, S. J., & Barnes, K. L. (1993). The reliability of ERP components in the auditory oddball paradigm. Psychophysiology, 30(5), 451–459. 10.1111/j.1469-8986.1993.tb02068.x

Shaw, M. E., Hämäläinen, M. S., & Gutschalk, A. (2013). How anatomical asymmetry of human auditory cortex can lead to a rightward bias in auditory evoked fields. NeuroImage, 74, 22–29. 10.1016/j.neuroimage.2013.02.002

Sielużycki, C., Matysiak, A., König, R., & Iskander, D. R. (2021). Reducing the Number of MEG/EEG Trials Needed for the Estimation of Brain Evoked Responses: A Bootstrap Approach. IEEE Transactions on Biomedical Engineering, 68(7), 2301–2312. 10.1109/ TBME.2021.3060495

Tadel, F., Baillet, S., Mosher, J. C., Pantazis, D., & Leahy, R. M. (2011). Brainstorm: A User-Friendly Application for MEG/EEG Analysis. Computational Intelligence and Neuroscience, 2011, 879716. 10.1155/2011/879716

Tiitinen, H., May, P., Reinikainen, K., & Näätänen, R. (1994). Attentive novelty detection in humans is governed by pre-attentive sensory memory. Nature, 372(6501), 90–92. 10.1038/372090a0

Tomana, E., Härtwich, N., Rozmarynowski, A., König, R., May, P. J. C., & Sielużycki, C. (2023). Optimising a computational model of human auditory cortex with an evolutionary algorithm. Hearing Research, 439, 108879. 10.1016/j.heares.2023.108879

v. Economo, C., & Horn, L. (1930). Über Windungsrelief, Maße und Rindenarchitektonik der Supratemporalfläche, ihre individuellen und ihre Seitenunterschiede. Zeitschrift für die gesamte Neurologie und Psychiatrie, 130(1), 678–757. 10.1007/ BF02865945

Wacongne, C., Changeux, J.-P., & Dehaene, S. (2012). A Neuronal Model of Predictive Coding Accounting for the Mismatch Negativity. Journal of Neuroscience, 32(11), 3665–3678. 10.1523/JNEUROSCI.5003-11.2012

Wang, A. L., Mouraux, A., Liang, M., & Iannetti, G. D. (2008). The Enhancement of the N1 Wave Elicited by Sensory Stimuli Presented at Very Short Inter-Stimulus Intervals Is a General Feature across Sensory Systems. PLOS ONE, 3(12), e3929. 10.1371/ journal.pone.0003929

Wernicke, C. (1874). Der aphasische Symptomencomplex: Eine psychologische Studie auf anatomischer Basis. Cohn & Weigert, Breslau

Willmore, B. D. B., & King, A. J. (2023). Adaptation in auditory processing. Physiological Reviews, 103(2), 1025–1058. 10.1152/physrev.00011.2022

Woldorff, M. G., Tempelmann, C., Fell, J., Tegeler, C., Gaschler-Markefski, B., Hinrichs, H., Heinz, H. J., & Scheich, H. (1999). Lateralized auditory spatial perception and the contralaterality of cortical processing as studied with functional magnetic resonance imaging and magnetoencephalography. Human Brain Mapping, 7(1), 49–66. 10.1002/(SICI)1097-0193(1999)7:1<49::AID-HBM5>3.0.CO;2-J

Zacharias, N., König, R., & Heil, P. (2012). Stimulation-history effects on the M100 revealed by its differential dependence on the stimulus onset interval. Psychophysiology. 49(7), 909–919 10.1111/j.1469-8986.2012.01370.x

Zatorre, R. J., Belin, P., & Penhune, V. B. (2002). Structure and function of auditory cortex: Music and speech. Trends in Cognitive Sciences, 6(1), 37–46. 10.1016/S1364-6613(00)01816-7

